# A quantitative analysis of the similarities and differences of the HIV/FIV gag genome

**DOI:** 10.1101/2022.01.09.475545

**Authors:** Charlotte Siu, Meredith Horn, Xiao-Wen Cheng

**Affiliations:** Miami University

**Keywords:** HIV, FIV, gag genome, gag proteins, FASTA, LALIGN, bioinformatics, homology, bit score, E value, percent identity

## Abstract

Among viruses, human immunodeficiency virus (HIV) presents the greatest challenge to humans. Here, we retrieved genome sequences from NCBI and were then run through LALIGN bioinformatics software to compute the E value, bit score, Waterman eggert score, and percent identity, which are four important indicators of how similar the sequences are. The E value was 3.1 x 10^-9, the percent identity was 54.4 percent, and the bit score was 51.9. It was also sensed that bases 1600 to 1990 in HIV and bases 800 to 910 in FIV have a higher than normal similarity. This reflects that while the DNA sequences of the gag region of both the HIV and FIV genomes are rather similar, it is unlikely that this similarity is due to random chance; therefore, there are a noticeable number of differences. A better understanding of the level of similarity and differences in the gag region of the genome sequence would facilitate our understanding of structural and cellular behavioral differences between FIV and HIV, and in the long term, it will provide new insights into the differences observed in previous studies or even facilitate the development of an effective HIV treatment.

## 2. Introduction

First discovered in 1983, HIV has infected approximately 80 million people worldwide so far according to data published by the WHO. In 2020, over 680,000 people died from the virus^1^. Despite its deadliness, there is still no effective and specific treatment for this virus. Three years after HIV was first sequenced, scientists were able to discover a similar virus in cats, feline immunodeficiency virus or FIV (Pederson et al, 1987)^2^. Like many viruses, both HIV and FIV have gag proteins (Coffin et al, 1997)^3^. The gag protein is known to play an important role in many stages of the replication cycle of a retrovirus. For example, they play an important role in viral assembly, interact with numerous host cell proteins, and regulate viral gene expression. They also provide the main driving force for virus intracellular trafficking and budding and are involved in pathogenicity (Mullers, 2013)^4^. Past studies have suggested that the DNA sequences in both viruses are similar, but it was not clear to what extent the gag genome similarity is^5^. When computed in the DNA analysis and alignment software FASTA, the E value and bit score are good indications of similarity between two sequences. The lower the E value is, the more similar the DNA sequences are, and the less likely this “match” in the DNA sequence is due to random chance. Generally, an E value below 0.01 is considered low. A bit score of 50 or above almost always indicates that the match between two DNA sequences is very significant and similar (Pearson, 2013).^6^ The percent identity is the percent of nucleotides that match exactly and is adjusted for the length of the DNA sequence. A percent identity of 50 percent or more would mean that a majority of the nucleotides match when adjusted for the length of the DNA sequences. Since previous studies have shown that HIV and FIV share similar pathogenesis and the gag protein coded by the gag genome plays an important role in pathogenesis (Friedman et al, 2006)^7^, it is hypothesized that the gag region of the FIV and HIV genome should be similar in sequence as defined by the aforementioned standards regarding the E value, bit score and percent identity.

## 3. Materials and Methods

### 3.1 Preparing the PCR templates

Before HIV and FIV DNA could be sequenced, the DNA samples needed to be amplified by PCR. This could increase the number of copies of the same DNA available for sequencing (Casbon et al, 2011).^8^

In this experiment, the master mix of the PCR consisted of 1 μL of big dye terminator (table 1), 1.5 μL of big dye dilution buffer (table 1), 0.5 μL of the primer (table 1), 4.5 μL of the gag DNA (table 1), and 5.5 μL of molecular grade water (table 1). This adds up to a total volume of 10 μL. The same recipe was used for both the HIV and FIV gags.

**Table 1.** The ingredients used to make the PCR master mix for both HIV and FIV gag and their respective quantities

| Ingredient | Quantity (µL) |
| --- | --- |
| Big dye terminator | 1 |
| Big dye dilution buffer | 1.5 |
| HIV/ FIV primer | 0.5 |
| HIV/ FIV gag DNA | 4.5 |
| Molecular grade water | 5.5 |

### 3.2 The PCR run

After preparing the master mix, we programmed the standard cycle sequencing protocol on the thermocycler. Step one of the cycles lasted for 1 minute at 96 °C, step 2 lasted 10 seconds at 96 °C, step 3 lasted 5 seconds at 50 °C, and step 4 lasted 4 minutes at 60 °C. The cycler was repeated for 35 times.

### 3.3 purification of the PCR products

After completing the PCR run, the master mix was transferred to a 1.5 mL Eppendorf tube, to which 1 μL of 1.5 M NaOAC/EDTA and 80 μL 95% ethanol were added. After that, the master mix was centrifuged for 15 minutes at 12,000 g. All supernatants were removed with a pipette, and 100 μL of 70% ethanol was added. The sample was stored at −20 °C.

### 3.4 the next generation sequencing machine

After the purification and PCR process, the sample was handed over to a technician to run on an ABI 3100 machine.

### 3.5 Checking the accuracy

The gag genome sequences received were “nucleotide blasted” on NCBI. The BLAST results showed that the sequences were accurate because they showed a 100% match to the HIV and FIV gag genome sequences in their records.

### 3.6 Bioinformatics software analysis

The HIV and FIV gag sequences obtained were entered into FASTA software, a software designed to compare DNA sequences. FASTA software was used to compute the E value, bit score and percent identity of the two sequences. The results were recorded and presented below.

## 4. Results

The E value of the overall sequences was 3.1×10^-9, with a bit score of 51.9. The percent identity of the overall sequences was 54.4% (Table 2). In addition, exactly aligning the nucleotide bases is shown in figure 1 below.

**Table 2.**
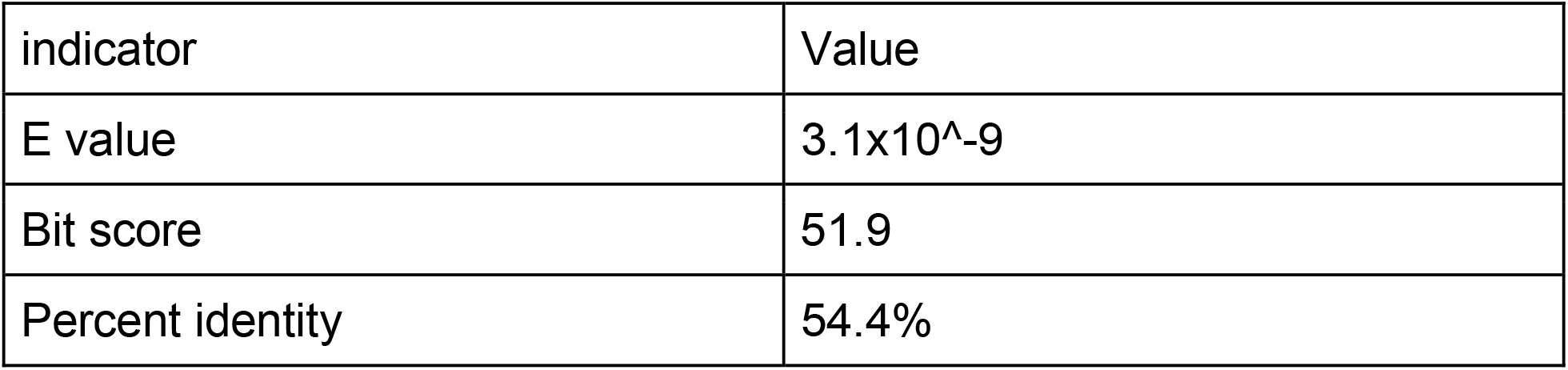
The LALIGN results of the HIV/ FIV gag genome

**Figure 1.**
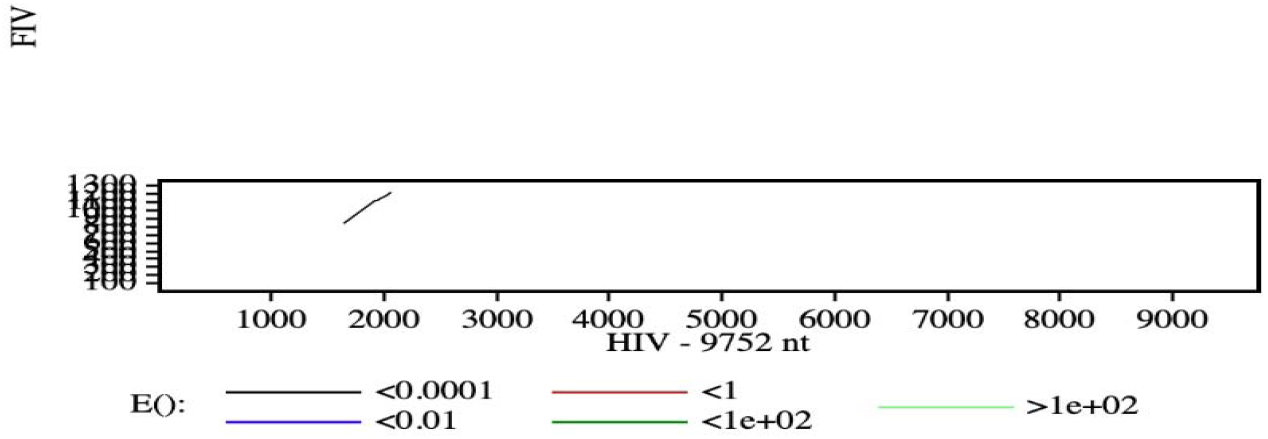
the software indicates that there is a specific region of the DNA in FIV and HIV that requires attention, since the E value is particularly low which means it is very similar and it is unlikely that it is by chance.

The results of the study support our hypothesis. The gag genomes of HIV and FIV, on a whole, are very similar. While they are generally similar, approximately 46.6 percent of the nucleotide bases do not exactly match.

In addition to calculating the E value, bit score and percent identity, the software alerted us that there is a region in the HIV/FIV genome that special attention is needed, since it shows a striking degree of similarity that can be said as one of the highest in the overall gag genome. It should be somewhere between the base pair 800 to 1,200 in FIV and 1,500 to 2,500 in HIV (Figure 1).

With the software’s alert in hand, a FASTA analysis was performed. Unlike LALIGN, the FASTA results show a more specific picture of where this “region of concern” is. The “region of concern” is base 1600 to 1990 in HIV and base 800 to 910 in FIV (Figure 2). The “region of concern” shows a 3.4×10^-10 E value and a 55.1 bit score (Figure 2). Both of these are higher than the overall gag genome.

**Figure 2.**
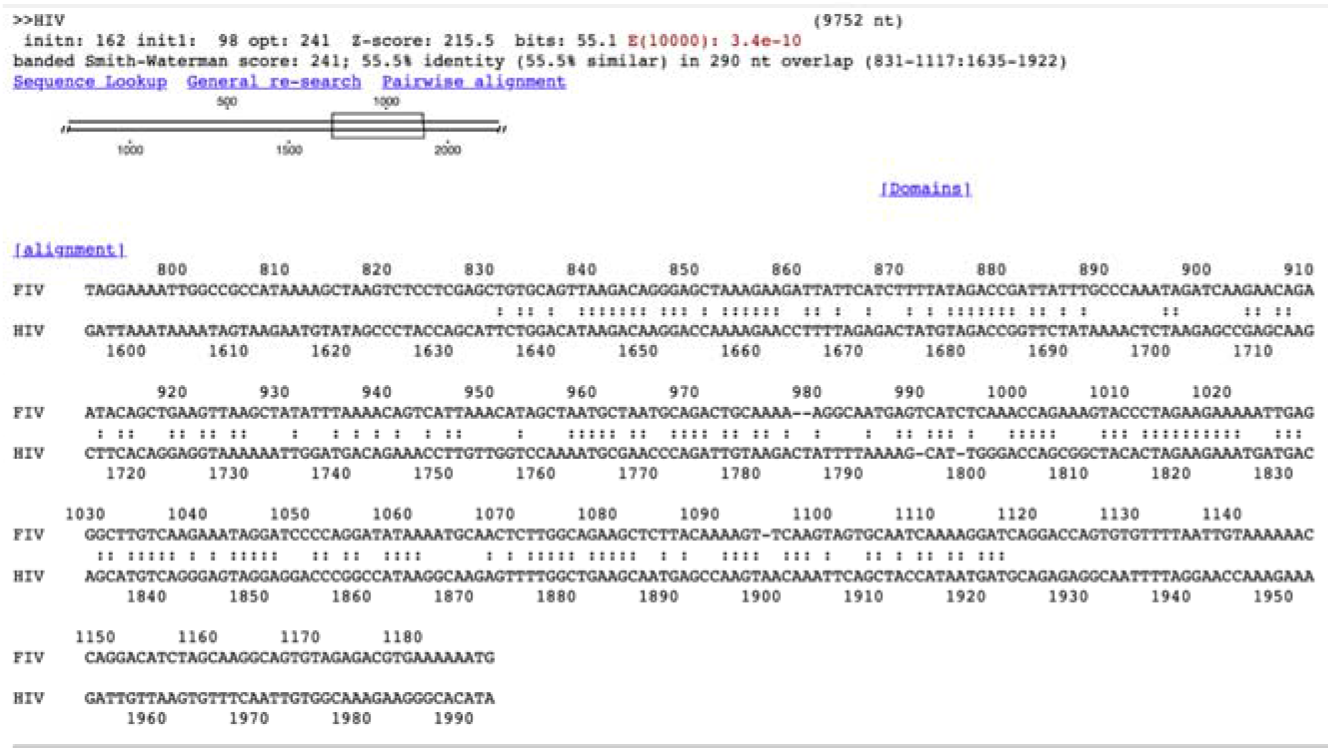
a FASTA analysis of the “region of concern”

## 5. Conclusions and discussions

Since DNA, RNA and protein are important parts of the central dogma, the answer to why these structural and cellular behavioral differences shown in gag proteins could lie within the gag genome itself. Building on what this research shows, subsequent studies could try to identify regions of the two gag genomes that show more differences than usual. What these regions code for will be of special interest to perform further research. In addition, what the aforementioned region codes for is something we previously did not know. Previous studies have concluded that FIV Gag is a nuclear shuttling protein that utilizes the CRM1 nuclear export pathway, while HIV-1 Gag is excluded from the nucleus (Kemler et al, 2012)^9^. The FIV matrix is not found as a trimer in the crystal structure. Existing research shows that there is little similarity in the amino acid sequence of the FIV and HIV protein matrix. However, there is evidence that the FIV capsid protein retains the same fundamental structure as the HIV protein (Gonzalez et al, 2018)^10^. Since proteins perform most enzymatic functions in all cells, the answer to why these structural and cellular behavioral differences could be of interest for further investigation. Conducting further studies to investigate the aforementioned questions or on the gag genome could provide new insights into why there are similarities and differences between HIV and FIV gags, as mentioned above, or even help us understand more about how the gag protein works in ways we have never understood before. As the gag protein is highly involved in virus replication, understanding gag more could help us develop effective HIV treatment or prevention in the long term.

## 6. Ethics approval

This study was approved by the Department of Microbiology at Miami University. All biosafety guidelines were followed.

## 7.2. Consent for publication

Not applicable.

## 7.3. Availability of data and materials

Data and materials are available upon request to the corresponding author.

## 9. Author contact information

Charlotte Siu

Meredith Horn

Xiao Wen Cheng

## Footnotes

1 https://www.who.int/data/gho/data/indicators/indicator-details/GHO/number-of-deaths-due-to-hiv-aids#:~:text=Situation%20and%20trends%3A%20680%2

2 https://ui.adsabs.harvard.edu/abs/1987Sci...235..790P/abstract

3 https://www.ncbi.nlm.nih.gov/books/NBK19464/

4 https://www.ncbi.nlm.nih.gov/pmc/articles/PMC3705263/

5 https://www.ncbi.nlm.nih.gov/pmc/articles/PMC5923500/

6 https://www.ncbi.nlm.nih.gov/pmc/articles/PMC3820096/

7 https://www.ncbi.nlm.nih.gov/pmc/articles/PMC7121254/

8 https://pubmed.ncbi.nlm.nih.gov/21490082/

9 https://pubmed.ncbi.nlm.nih.gov/22623802/

10 https://www.ncbi.nlm.nih.gov/pmc/articles/PMC5977254/

